# Nanopore sequencing significantly improves genome assembly of the eukaryotic protozoan parasite *Trypanosoma cruzi*

**DOI:** 10.1101/489534

**Authors:** Florencia Díaz-Viraqué, Sebastian Pita, Gonzalo Greif, Rita de Cássia Moreira de Souza, Gregorio Iraola, Carlos Robello

**Author notes:** Author for Correspondence: Carlos Robello, Laboratorio de Interacciones Hospedero-Patógeno, Institut Pasteur de Montevideo, Uruguay. Gregorio Iraola, Laboratorio de Genómica Microbiana, Institut Pasteur de Montevideo, Uruguay.

## Abstract

Chagas disease was described by Carlos Chagas, who first identified the parasite *Trypanosoma cruzi* from a two-year-old girl called Berenice. Many *T. cruzi* sequencing projects based on short reads have demonstrated that genome assembly and downstream comparative analyses are extremely challenging in this species, given that half of its genome is composed of repetitive sequences. Here, we report *de novo* assemblies, annotation and comparative analyses of the Berenice strain using a combination of Illumina short reads and MinION long reads. Our work demonstrates that Nanopore sequencing improves *T. cruzi* assembly contiguity and increases the assembly size in ~16 Mb. Specifically, we found that assembly improvement also refines the completeness of coding regions for both single copy genes and repetitive transposable elements. Beyond its historical and epidemiological importance, Berenice constitutes a fundamental resource since it now represents the best-quality assembly available for TcII, a highly prevalent lineage causing human infections in South America. The availability of Berenice genome expands the known genetic diversity of *T. cruzi* and facilitates more comprehensive evolutionary inferences. Our work represents the first report of Nanopore technology used to resolve complex protozoan genomes, supporting its subsequent application for improving trypanosomatid and other highly repetitive genomes.

## Introduction

The MinION (Oxford Nanopore Technologies) is an instrument that fits in the palm of a hand and can be plugged in a laptop computer allowing single-molecule, real-time DNA sequencing with unprecedented speed and portability. Since this instrument directly reads individual DNA fragments without the need for amplification steps during library preparation, it has the capacity of producing very long reads. The availability of this kind of sequencing data is relevant when assembling genomes that are rich in repetitive elements, because long reads allow to span entire tandems of repeats and anchor them to uniquely occurring segments of the genome, resolving these complex regions and improving contiguity. However, the still high error rates of MinION demands considerable amounts of data and intensive computation to build entire genomes just using long reads. Conversely, hybrid strategies that combine error-prone long reads with much more accurate Illumina short reads represent a more convenient approach for enhancing genome completeness. Indeed, several organisms ranging from bacteria (Wick et al. 2017) to vertebrates (Tan et al. 2018) have been recently sequenced using a combination of Nanopore and Illumina reads. However, this strategy has not been implemented so far to resolve protozoan genomes.

*Trypanosoma cruzi* is a protozoan parasite belonging to the order *Kinetoplastida* that causes Chagas disease (CD), also known as American Trypanosomiasis, a neglected parasitic disease that affects 6-7 million people worldwide and is transmitted to humans and animals mainly by Triatomine insect vectors (Deane 1964; WHO, 2017). CD recently emerged in non-endemic regions such as Western Europe, Australia, Japan, Canada and the United States due to widespread immigration, however its highest incidence is observed in Latin American countries where the parasite is endemic (Rassi et al. 2010). Indeed, CD was first diagnosed in Brazil more than one century ago by Carlos Chagas when he examined the two year old girl Berenice Soares (Chagas 1909), who developed the asymptomatic form of the disease (de Lana et al. 1996). The archetypal *T. cruzi* strain originally isolated from this case (Salgado et al. 1962) represents the oldest known record for this pathogenic parasite, and own invaluable historical, cultural and epidemiological importance. The Berenice strain has been characterized in many aspects, but has not been whole-genome sequenced by any technology so far.

Here, we report the whole-genome sequence, annotation and comparative analysis of the Berenice strain isolated by Salgado et al. (1962) using a combination of Illumina short reads and MinION long reads, providing a useful genetic resource for the community working with parasite genomes. Importantly, we demonstrate that a single MinION run based on a straightforward 10-minute library preparation protocol allows a 67-fold increase in genome contiguity and improves genome completeness by 28% when compared with short-read-only assemblies. Our results show that hybrid assembly strategies using MinION are effective when dealing with complex protozoan genomes like *T. cruzi*.

## Results

### Nanopore sequencing improves *T. cruzi* assembly contiguity and size

We whole-genome sequenced *T. cruzi* strain Berenice using Illumina 150 bp pair-end short reads and Nanopore 1D long reads (Table 1). Then, we produced two genome assemblies, one just using the short reads from Illumina (hereinafter referred as the Illumina assembly) and the other by combining Illumina short reads with Nanopore long reads (hereinafter referred as the hybrid assembly). Figure 1A shows a 46-fold improvement in median scaffold size in the hybrid assembly. This improvement is also evident by a 51-fold decrease in scaffold number (from ~47,000 scaffolds with a maximum length of ~26 Kb in the Illumina assembly to ~900 scaffolds with a maximum length of ~1 Mb in the hybrid assembly), and a ~16 Mb increase in assembly size product of improved resolution of repeated regions (Table 1). Also, the cumulative hybrid assembly size is kept practically unchanged around ~40 Mb when considering scaffolds of increasing size, evidencing insignificant contribution of small scaffolds to the whole assembly. On the contrary, the cumulative size of the Illumina assembly rapidly tends to zero when considering longer scaffolds evidencing an extremely fragmented assembly (Fig. 1B).

**Figure 1.**
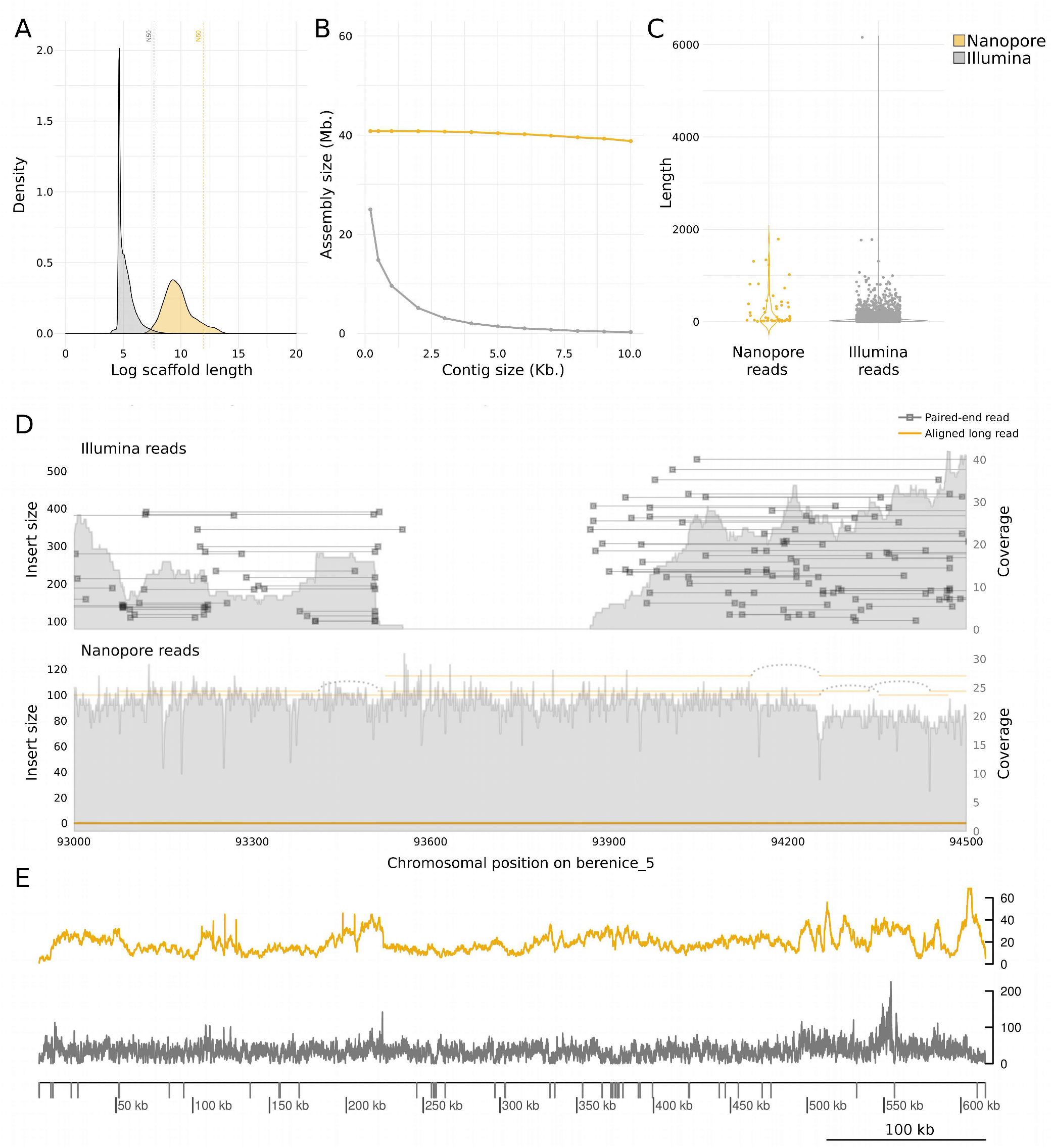
*Nanopore sequencing improves T. cruzi* assembly contiguity and size. A) Scaffolds length distribution. N50 of the hybrid genome assembly: 156193. N50 of Illumina assembly: 2127. B) Cumulative assemblies size. C) Coverage zero regions (no read alignment in at least 6 consecutive positions) observed when Nanopore or Illumina reads were aligned to the hybrid assembly in order to assess the contribution of both technologies to the assembly contiguity D) Sequencing coverage and insert size from 93 Kb to 94.5 Kb positions of scaffold berenice_5 from hybrid assembly are plotted. E) Per-base genome coverage of scaffold 4 of hybrid assembly. Coverage zero regions are plotted as gray bars over the exe and all were observed when Illumina reads were aligned to hybrid assembly.

**Table 1.** Summary of genome assemblies and annotation.

|  | Hybrid genome assembly | Illumina genome assembly |
| --- | --- | --- |
| **Assembly features** |  |  |
| Number of contigs | 923 | 46,821 |
| Largest contig | 926,516 | 26,836 |
| Size (bp) | 40,801,262 | 25,004,252 |
| GC (%) | 51.20 | 48.67 |
| N50 | 156,193 | 659 |
| N75 | 40,889 | 333 |
| Number of contigs (>= 50,000 pb) | 160 | 0 |
| **Annotation** |  |  |
| Coding genes | 14,032 | 4,282 |
| Non-coding genes | 456 | 107 |
| tRNA | 58 | 47 |
| tRNA-Sec | 1 | 0 |
| 5S rRNA | 20 | 1 |
| 5.8S rRNA | 7 | 1 |
| SSU | 10 | 3 |
| LSU | 10 | 7 |
| snoRNA | 345 | 46 |
| snRNA | 5 | 2 |
| Transposable elements | 388 | 4 |
| CZAR | 27 | 0 |
| L1Tc | 38 | 0 |
| VIPER | 50 | 0 |
| NARTc | 54 | 4 |
| SIRE | 80 | 0 |
| TcTREZO | 139 | 0 |
| **Completeness** |  |  |
| Complete BUSCO eukaryote | 206 (68 %) | 147 (48.5 %) |
| Fragmented BUSCO eukaryote | 19 (6.3 %) | 62 (20.5 %) |
| Missing BUSCO eukaryote | 78 (25.7 %) | 94 (31 %) |
| Complete BUSCO protis | 121 (56.3 %) | 61 (28.4 %) |
| Fragmented BUSCO protis | 1 (0.5 %) | 1 (0.5 %) |
| Missing BUSCO protis | 93 (43.3 %) | 153 (71.2 %) |

### Nanopore reads close Illumina assembly gaps

To evaluate the contribution of Illumina and Nanopore data to close gaps, we separately aligned both types of reads to the hybrid assembly. The longest region where the coverage is zero (no read alignment in at least 6 consecutive positions) spanned 6,156 bp with Illumina reads while it decreased to 1,787 bp with Nanopore reads. Additionally, assembly regions of coverage zero were much more abundant when aligning Illumina reads (n=3624) than when aligning Nanopore reads (n=54) (Fig. 1C). One of these regions is represented in Fig. 1D, where Nanopore reads uninterruptedly cover this genomic segment with a smooth depth of ~20x while Illumina reads fail to resolve an intrinsic region where coverage falls to zero, causing the break of contiguity in the assembly.

### Nanopore reads improve completeness of coding regions

To assess whether assembly improvement also refines the completeness of coding regions, we first annotated protein-coding genes and non-coding RNA genes. We obtained a 3-fold increase in the recovery of protein-coding genes, non-coding RNA genes and transposable elements from the hybrid assembly in comparison with the Illumina assembly (Table 1). Additionally, we tested completeness by attempting the recovery of conserved single-copy genes from both assemblies. Out of a database containing more than 215 single-copy protozoan orthologs, ~57% were fully recovered from the hybrid assembly while only ~29% were recovered from the Illumina assembly. Also, when using a more general database containing over 303 single-copy orthologs conserved across eukaryotic organisms, 68% of these genes were recovered from the hybrid assembly while 48.5% from the Illumina assembly. Together, this demonstrates that Nanopore sequencing helps to mitigate the underestimation of both unique and repetitive coding regions of the genome.

### Berenice belongs to TcII and is phylogenetically close to Esmeraldo

*T. cruzi* strains are traditionally classified in discrete typing units (DTUs) TcI-TcVI based on molecular and phylogenetic analyses. Based on this, Berenice belongs to TcII (Zingales et al. 2009). To test this, we performed a phylogenetic analysis including several available *T. cruzi* genomes and Berenice using L1Tc sequences, previously defines as an accurate molecular clock (Berná et al., 2018). The resulting tree showed three major lineages (Fig. 2), one comprising sequences from Dm28c and Silvio that defined TcI, other conformed mainly of sequences from TCC and Non-Esmeraldo that defined TcIII, and the remaining composed by Berenice, TCC and Esmeraldo, confirming that Berenice belongs to TcII.

**Figure 2.**
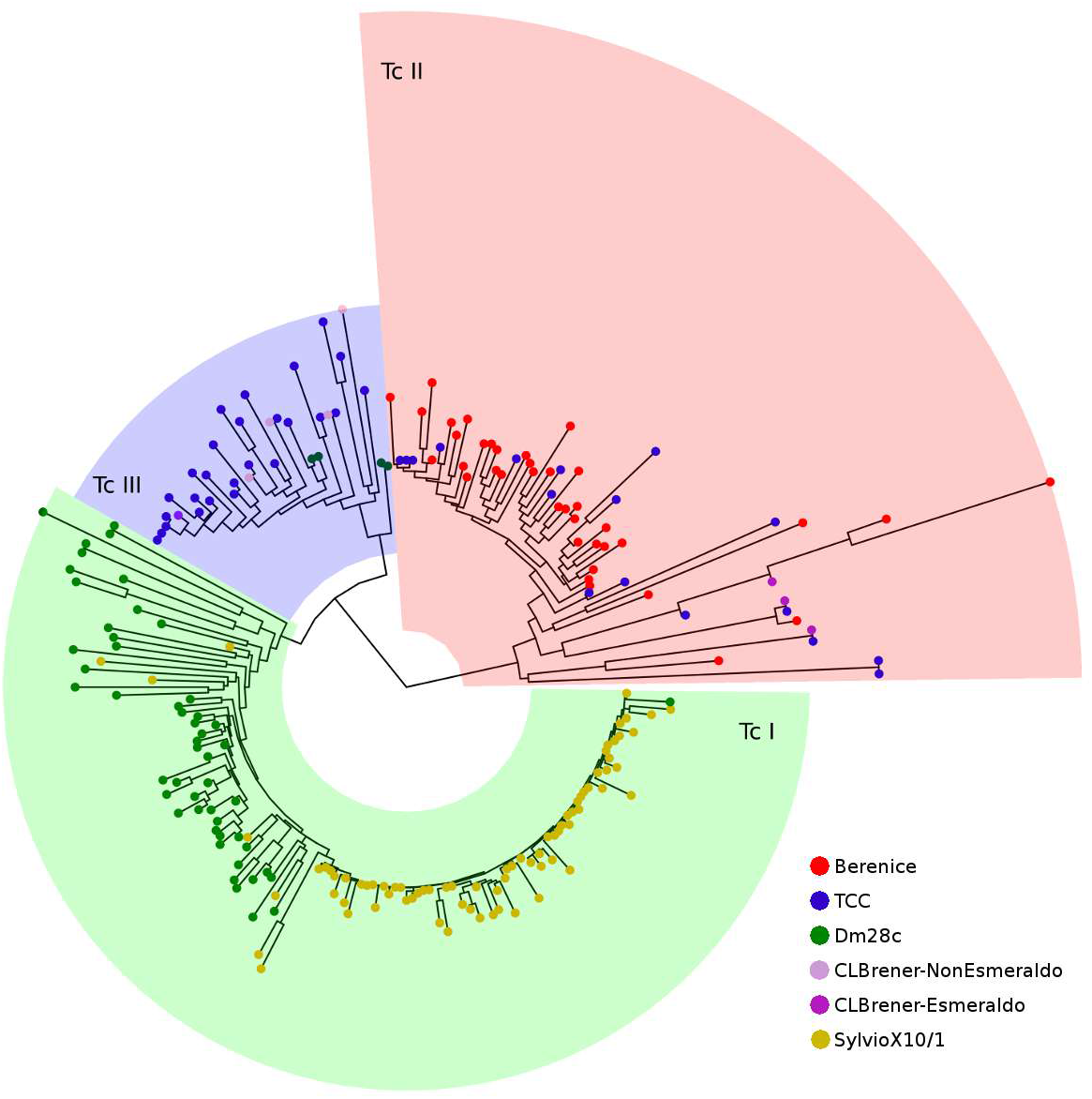
Evolutionary relationships of *T. cruzi* strains. Maximum-likelihood phylogeny constructed with full L1Tc sequences recovered from six *T. cruzi* genomes.

## Discussion

*Trypanosoma cruzi* is the causative agent of Chagas disease, an important neglected tropical disease that affects about 6-7 million people worldwide (WHO, 2017). Here, we report the complete genome sequence of Berenice, the first *T. cruzi* strain isolated from a patient (Chagas 1909). This represents the first trypanosomatid parasite genome generated using a hybrid assembly strategy by combining Illumina short reads and Nanopore long reads.

Even though trypanosomatid genomes are small, their assembly and annotation have been challenging due to the abundance of repetitive sequences including the 195bp satellite, tandem repeats, and multigene families (El-Sayed et al. 2005a; Berná et al. 2018; Pita et al. 2018). In fact, when “tritryp” genomes were sequenced in 2005, *T. cruzi* genome assembly remained highly fragmented (El-Sayed et al. 2005b), hampering highly precise comparative genomics. However, the recent advent of long-read sequencing technologies such as PacBio and Oxford Nanopore is allowing us to overcome these limitations.

Long-read sequencing using PacBio has been proven useful to improve the quality of *T. cruzi* genome assemblies (Berná et al. 2018; Callejas-Hernández et al. 2018), however, the innovative Nanopore technology has been not implemented to sequence trypanosomatid genomes so far despite presenting several comparative advantages over PacBio. Nanopore is cheaper, easy to use in any laboratory, requires less amount of genomic DNA and sequencing yield can be monitored in real-time. Additionally, Nanopore offers countless possibilities for library preparation including quick, straightforward protocols. Indeed, here we show that a 10-minute library preparation protocol followed by 12 hours of Nanopore 1D sequencing significantly improves assembly contiguity and annotation, demonstrating the usefulness of this technology to resolve highly complex parasite genomes.

Beyond its historical, cultural and epidemiological relevance for being an isolate from the first clinical case of Chagas disease, Berenice strain was chosen in order to increase the phylogenetic representativeness of genomes resolved by long-read sequencing. To date, three *T. cruzi* strains have been assembled with long reads: strain Dm28c belonging to TcI, strain Bug2148 belonging to TcV and strain TCC belonging to TcVI. Now, Berenice represents a fundamental resource since it becomes the best-quality assembly available for a member of TcII (Table 2), contributing to expand the known genetic diversity of *T. cruzi* and facilitating the production of more comprehensive evolutionary inferences. It was almost two decades ago when the presence of three major groups of *T. cruzi* strains was described for the first time (Robello et al., 2000). Indeed, TcI, TcII and TcIII are homozygous lineages, being TcII and TcIII proposed as the putative parents for the remaining hybrid lineages TcV and TcVI (de Freitas et al. 2006; Zingales et al. 2012). Concordantly, our phylogenetic analysis places Berenice close to TCC and CLBrener Esmeraldo-like, both from TcVI lineage.

**Table 2.** Comparison of the *T. cruzi* genomes assembled with long read.

|  | Sequencing method | Size (Mb) | GC (%) | Number of contigs | N50 | Coverage | BUSCO eukaryota |
| --- | --- | --- | --- | --- | --- | --- | --- |
| Dm28c (TcI) | PacBio | 53.16 | 51.6 | 599 | 317,638 | 76 X | 202 |
| Berenice (TcII) | Illumina Nanopore | 40.80 | 51.2 | 923 | 156,193 | 41 X Illumina<br>28 X Nanopore | 206 |
| Bug2148 (TcV) | PacBio | 55.22 | 51.63 | 934 | 196,760 | 68 X | 208 |
| TCC (TcVI) | PacBio | 86.77 | 51.7 | 1142 | 265,169 | 60 X | 210 |

Here, we used a combination of Illumina and Oxford Nanopore reads to provide the most complete genome assembly of a TcII *T. cruzi* strain and is the first report of Nanopore reads used for trypanosomatids. We compared the assembly continuity and completeness obtained with the most simple library preparation kit of Nanopore with the assembly obtained only using Illumina reads and we obtained a highly improved assembly, similar to the ones obtained using PacBio reads. Even though the coverage and libraries preparation can be optimized, we demonstrate that Oxford Nanopore can be a very valuable technology to improve highly repetitive genomes such as trypanosomatids. This approach has several advantages and can be carried out in every laboratory without any previous training in sequencing, contributing to facilitate the enlargement of genomic resources for protozoan pathogens.

## Methods

### Library preparation, genome sequencing and assembly

Genomic libraries were prepared with the Nextera^®^ XT Library Prep Kit (Illumina, 15032354) and Rapid Sequencing Kit (Nanopore, SQK-RAD004). Illumina and Nanopore libraries were sequenced in MiSeq and MinION platforms, producing 12,589,973 paired-end short reads and 265,221 long reads, respectively. Integrity of Illumina libraries were analyzed using 2100 Bioanalyzer (Agilent) and quantified using Qubit™ dsDNA HS Assay Kit. Berenice genome assembly was performed using Illumina reads (Illumina genome assembly) and mixing Illumina and Nanopore reads (Hybrid genome assembly) with MaSuRCA using default parameters (Zimin et al. 2013, 2017).

### Comparison of genome assemblies

For genome assembly comparisons, Illumina and Nanopore reads were aligned to Berenice genome assembled with both reads using minimap2 v2.10-r784 (Li 2018) with default parameters. Per-base genome coverage was calculated using bedtools v2.26.0 (Quinlan and Hall 2010) and samplot (Belyeu et al. 2018) was used for rendering the sequencing coverage in specific genomic regions. Completeness of genome coding regions was assessed using BUSCO v3.0.2 (Simão et al. 2015) with the eukaryotic and protist linages databases.

### Genome annotation

In order to annotate the coding sequences, the annotated proteins of 41 protozoan parasites genomes were obtained from TriTrypDB release 38 (http://tritrypdb.org/). Otherwise, all open reading frames longer than 150 amino acids were retrieved between start and stop codon using getorf from the EMBOSS suite (Rice et al. 2000) in both assemblies. Homologous genes were recovered using BLAST+ blastp (Camacho et al. 2009), with alignment coverage >80%, identity percentage > 80% and an e-value threshold of 1e-10. Rfam release 13 (Nawrocki et al. 2015) and Infernal v1.1.1 (Nawrocki and Eddy 2013) were used for the annotation of non-coding genes as it was previously described (Kalvari et al. 2018). For tRNAs, tRNAscan-SE v.1.3.1 (Lowe and Chan 2016) was used with the euakyotic model. Transposable elements were annotated using BLAST+ blastn (Camacho et al. 2009) and tandem repeats were annotated using Tandem Repeat Finder v4.09 (Benson 1999).

### Phylogenetic analysis

Complete nucleotide sequences of L1Tc transposable elements were used to perform phylogenetic analyses. Sequences retrieved from six genomes were aligned using MAFFT v7.310 (Katoh and Standley 2013) with the L-ins-i option. A Maximum-Likelihood phylogenetic tree was reconstructed using PhyML v20120412 (Guindon et al. 2010) using the best-fitted model GTR selected with ModelGenerator v0.85 (Benson 1999).

## Data Access

Sequencing data generated in this work has been deposited at the NCBI repository under the BioProject accession PRJNA498808.

## Acknowledgments

This work was funded by Agencia Nacional de Investigación e Innovación (ANII) DCI-ALA/2011/023-502, ‘Contrato de apoyo a las políticas de innovación y cohesión territorial’, Fondo para la Convergencia Estructural del Mercado Común del Sur (FOCEM) 03/11, and by Research Council United Kingdom Grand Challenges Research Funder ‘A Global Network for Neglected Tropical Diseases’ grant number MR/P027989/1. SP, GI, GG and CR are members of the “Sistema Nacional de Investigadores (ANII)”; FDV has an ANII doctoral fellowship No. POS_NAC_2016_1_129916.

## Author contributions

CR and GI conceived the idea. RCM processed and prepared samples. FDV and GG prepared libraries and performed Illumina and Nanopore sequencing. FDV, GI, and SP analyzed the data. FDV, GI and CR wrote the manuscript. All authors approved the final version of the manuscript.

